# Analysis of model organism viability through an interspecies pathway comparison pipeline using the dynamic impact approach

**DOI:** 10.1101/2019.12.18.448985

**Authors:** Austin Nguyen, Massimo Bionaz

## Abstract

**Background:** Computational biologists investigate gene expression time-series data using estimation, clustering, alignment, and enrichment methods to make biological sense of the data and provide compelling visualization. While there is an abundance of microarray and RNA-seq data available, interpreting the data while capturing the dynamism of a time-course experiment remains a difficult challenge. Advancements in RNA-seq technologies have allowed us to collect extensive profiles of diverse developmental processes but also requires additional methods for analysis and data integration to capture the increased dynamism. An approach that can both capture the dynamism and direction of change in a time-course experiment in a holistic manner and simultaneously identify which biological pathways are significantly altered is necessary for the interpretation of systems biology data. In addition, there is a need for a method to evaluate the viability of model organisms across different treatments and conditions. By comparing effects of a specific treatment (e.g., a drug) on the target pathway between multiple species and determining pathways with a similar response to biological cues between organisms, we can determine the best animal model for that treatment for future studies.

**Methods:** Here, we present Dynamic Impact Approach with Normalization (DIA-norm), a dynamic pathway analysis tool for the analysis of time-course data without unsupervised dimensionality reduction. We analyzed five datasets of mesenchymal stem cells retrieved from the Gene Expression Omnibus data repository (3 human, 1 mouse cell line, 1 pig) which were differentiated *in vitro* towards adipogenesis. In the first step, DIA-norm calculated an impact and flux score for each biological term using *p*-value and fold change. In the second step, these scores were normalized and interpolated using cubic spline. Cross-correlation was then performed between all the data sets with r≥0.6 as a benchmark for high correlation as r = 0.7 is the limit of experimental reproducibility.

**Results:** DIA-norm predicted that the pig was a better model for humans than a mouse for the study of adipogenesis. The pig model had a higher number of correlating pathways with humans (64.5 to 30.5) and higher average correlation (r = 0.51 vs r = 0.46) as compared to mouse model vs human. While not a definitive conclusion, the results are in accordance with prior phylogenetic and disease studies in which pigs are a good model for studying humans, specifically regarding obesity. In addition, DIA-norm identified a larger number of biologically important pathways (approximately 2x number of pathways) versus a comparable enrichment analysis tool, DAVID. DIA-norm also identified some possible pathways of interests for adipogenesis, namely, nitrogen metabolism (r = 0.86), where there is little to no existing literature.

**Conclusion:** DIA-norm captured 80+% of biological important pathways and achieved high pathway correlation between species for the vast majority of important adipogenesis pathways. DIA-norm can be used for both time-series pathway analysis and the determination of a model organism. Our findings indicate that DIA-norm can be used to study the effect of any treatment, including drugs, on specific pathways between multiple species to determine the best animal model for that treatment for future studies. The reliability of DIA-norm to provide biological insights compared to enrichment approach tools has been demonstrated in the selected transcriptomic studies by identifying a higher number of total and biologically relevant pathways. DIA-norm’s final advantage was its easily interpretable graphical outputs that aid in visualizing dynamic changes in expression.

## Introduction

Computational biologists investigate time-series gene expression data using estimation, clustering, alignment, and enrichment methods to make biological sense of the data and provide compelling visualization. While there is an abundance of microarray and RNA-seq data available, interpreting the data while capturing the dynamism of a time-course experiment remains a difficult challenge (Androulakis et al., 2007). The fundamental change from the analysis of a single gene’s influence on a condition to a pathway’s effect on a condition allows for the capturing of a holistic view of the results of a biological process. This significantly reduces the influence of a small change in single-gene expression. However, capturing too broad of a view (e.g. unsupervised reduction) can result in loss of variation, something that is especially important given the rise of personalized medicine (Nair, 2010). Large differences in responses to small biological changes are exhibited in transcriptomic studies in many species, including humans (Xu et al., 2011). Because of this, the subject variation, which causes larger distribution in values and is often indistinguishable from real biological changes, can make it more difficult to see effects. Advancements in RNA-seq technologies have allowed us to collect extensive profiling of diverse developmental processes but also require additional methods for analysis and data integration to capture the increased dynamism (Spies and Ciaudo, 2015). An approach that can both capture the dynamism and direction of change in a time-course experiment in a holistic manner and simultaneously identify which biological pathways are significantly altered is necessary for the interpretation of systems biology data (Martini et al., 2014).

Current approaches to analyzing transcriptomic data often involve clustering, enrichment analysis, and principal component analysis (PCA). Clustering, specifically K-means clustering, has mathematically been shown to be linked to PCA (Kriegel et al., 2008). Principal component analysis is best used for exploratory data analysis or predictive models. In addition, as it is the highest level of simplification of an eigenvector multivariate analysis, which is thought of as finding the most important variants in a data set, there is an inherent loss of variation during formation of an XX^T^ covariance matrix, which can cause loss of precision (e.g. a dataset with similar properties to a Lauchli Matrix) (Le et al., 2017). This decomposition often effectively reduces the complexities of a data set into a small set of high-variance genes. Although scientist study the first two to four principal components in most studies, it has been demonstrated that there are more than three to four dimensions in large gene expression datasets, in which PCA would not capture significant biological variability (Lenz et al., 2016). While it is currently impossible to capture and interpret complete biological variation, reliance on a clustering or principal component analysis method may not be optimal for biologically relevant interpretation of overall effects in time course experiments.

With the rise of gene-set hypothesis testing in systems biology, enrichment analysis methods have become the most popular style of analysis and many attempts to improve the original GSEA model have been undertaken (Tarca et al., 2013). The main limitation of enrichment or overrepresented approaches is their dependency on the background gene list (Huang et al., 2009a). Dependency on the background gene list arises when enrichment scores for GSEA calculated between gene sets of higher rank are affected by the presence (or absence) of a lower rank set (Damian and Gorfine, 2004). As the enrichment scores result in the ranking of gene set lists, the final result is impacted by the selected background. Enrichment approaches assume that correlation between a biological pathway/function and the genes associated with it suggests involvement of the pathway in the biological response or disease that is being profiled (Irizarry et al., 2009). This points to an enrichment or a higher proportion of differentially expressed genes over a random selection of genes in the background. While statistically logical, inter-gene correlation can affect the *p*-value of enrichment and increase the false discovery rate (Huang et al., 2009b). In addition, multiple comparisons oftentimes reduce the gene list to a very small number of extremely influential genes, making results very narrow and holistic interpretation difficult.

This reduction of the gene list also makes it difficult to compare conclusions between data sets, including data obtained from different individuals and species. Because the resulting gene set after analysis is small, individual gene differences (e.g. *p*-value or fold change as in the case of GSEE) produces a disproportionately large effect and makes it difficult to draw meaningful comparisons (Huang et al., 2015). Another drawback would be experiments with large effects on the transcriptome; for example, the process of adipogenesis (Bionaz et al., 2015). There will be many differentially expressed genes but due to the comparisons with the pathways that both contain genes in the random background and differentially expressed genes, it will result in few enriched pathways. In comparison, a condition with a small effect on expression of genes, i.e., with few differentially expressed genes, would have more likelihood of having genes that are not distributed in pathways as the genes in the background, resulting in a larger number of enriched pathways (Bionaz et al., 2012).

Thus, enrichment and clustering methods do not provide an adequate solution to time-course analyses. Tools such as BiNGO (Maere et al., 2005), Ontologizer (Bauer et al., 2008), gProfiler (Reimand et al., 2007), and Babelomics (Alonso et al., 2015) can be used to perform time-course analyses but are insufficient for a biologist to analyze time-series data without repeated analyses for every time point; this would create a number of distinct results and it would be difficult to easily compare between time points. We conclude that biologists and bioinformaticians need an uncomplicated tool that can provide visualization of the biological effect of genes significantly affected by the studied condition, the usage and interpretation of a large gene set, and the ability to easily compare results from time-series data.

There is also a need for a method to evaluate the viability of model organisms across different treatments and conditions (Reed et al., 2017). The quantitative measurement for the evolutionary conservedness by linear analysis of genes related to particular pathways can provide some indication about conservation of the response; however, this can be only demonstrated by subjecting the pathway to particular cues and determining the similarity in the overall response (Tipney and Hunter, 2010). Determining the best model organism for a study is not a trivial task; for example, zebrafish are commonly used for studying development due to their embryos’ transparency but comprehensive gene expression comparisons between zebrafish and humans during development have not yet been performed (Howe et al., 2017). By comparing effects of a specific treatment (e.g., a drug) on the target pathway between multiple species and determining pathways with a similar response to biological cues between organisms, we can determine the best animal model for that treatment for future studies.

*In vitro* adipogenesis has been widely studied due to its role on adiposity, and thus obesity by association (Spiegelman and Flier, 1996). For this reason, molecular regulation of adipogenesis and its key bioactive molecules are relatively well known. Adipogenesis can be divided into three stages: early (1-6 days), middle (6-11 days), and late stages (11+ days) (Moreno-Navarrete and Fernández-Real, 2012; Xue et al., 1996). Different pathways are up/downregulated at different stages of adipogenesis (Pei et al., 2011). Because of its distinct stages, the availability of data from several species, and the well-studied biological characterization, *in vitro* adipogenesis is ideal for testing the validity of the DIA-norm.

In this study, we propose a refined method of analysis, DIA-norm, which uses the previously developed algorithm denominated “Dynamic Impact Approach” or DIA (Bionaz et al., 2012). DIA was specifically developed to analyze time-course experiments without unsupervised dimensionality reduction. The objectives of the present work were: 1) to create a web-based online DIA tool for the analysis of time-course experiments; 2) to improve DIA for interpretation of data by adding z-scores, normalization, interpolation, and time-lagged correlation analysis that allows users to identify most important pathways (i.e., DIA-norm); 3) to demonstrate the validity of DIA-norm on capturing biological relevant pathways related to a well-define biological function; 4) to assess the feasibility of DIA-norm to determine the best animal model for the study of *in vitro* adipogenesis. Proof of concept was demonstrated through the analysis of adipogenesis of preadipocyte cells in three species (human, mouse, and pig).

## Methods

### Step 1: Experimental data retrieval and processing

Datasets were retrieved from the Gene Expression Omnibus (GEO) dataset repository (https://www.ncbi.nlm.nih.gov/gds) for human (GSE37836, GSE61302, GSE77532), mouse (GSE2192), and pig (GSE25854). Downloaded data were annotated using annotation present in GEO and statistical analysis was performed using JMP Genomics (v8, SAS Institute, USA) with time as main effect and subjects as random effects using a general linear model with false discovery rate correction (FDR) using Benjamini-Hochberg correction for the overall time effect (Benjamini and Hochberg, 1995). For each gene the following information was obtained from JMP Genomics for each time point comparison: FDR, fold change and associated *p*-value.

For human sample 1 (Satish et al., 2015), adipose stem cells (ASC) were isolated from human patients and kept in DMEM supplemented with 10% FBS. Adipogenesis was induced with a cocktail composed of 1% penicillin/streptomycin, 0.5 % Fungizone, and 0.001 % dexamethasone. Cells were harvested at 0, 7, and 21 days of differentiation. Human sample 2 (Ambele et al., 2016) had ASCs isolated from liposuction patients. The ASCs were kept in DMEM supplemented with 10% FBS and 2% insulin. Adipogenesis was induced by an adipogenic cocktail composed of 1.0 μM dexamethasone, 0.5 mM isobutylmethylxanthine, 10 μM insulin, and 200 μM indomethacin. Cells were harvested at 0, 7, 14, and 21 days of differentiation. Human sample 3 (Hasegawa et al., 2013) used bone marrow stem cells purchased from RIKEN BRC and Takara Bioscience. The ASCs were maintained in MEM-α supplemented with 10% FBS plus adipogenic induction medium and adipogenic induction SingleQuots. Cells were harvested at 0, 0.5, 1, 2, 4, 6, 9, 12, and 15 days of differentiation. The mouse sample (Akerblad et al., 2005) used NIH-3T3 fibroblasts and induced adipogenesis using DMEM supplemented with 10% FBS plus an adipogenic differentiation medium composed of 1 μM dexamethasone, 0.5 mM isobutylmethylxanthine, 1 μg/ml insulin, and 0.5 μM darglitazone. Cells were harvested at 0, 2, 4, and 10 days of differentiation. The pig sample (Monaco et al., 2012) harvested ASC from back fat and cultured in high glucose DMEM supplemented with 10% FBS plus an adipogenic differentiation medium composed of 1.0 μM dexamethasone, 0.5 mM isobutylmethylxanthine, 10 μM insulin, and 200 μM indomethacin. Cells were harvested at 2, 7, and 21 days of differentiation.

To convert from the JMP Genomics output file to the DIA-norm input file, these columns are extracted: gene ID, PtF_T3_Time (FDR value for the overall time effect), Diff of Time columns with comparisons with time 0 (which is provided a the difference between log2 expression of the time point 0 and the log2 expression value of the time point of interest), and associated −log10 *p*-value columns. The fold changes were then obtained by elevating the number to exponential of 2, and converted to the 0 time-point as the denominator by inversing the value. The *p*-values were converted to normal *p*-value as 10 exponential of the −log10 *p*-value.

### Step 2: Pathway class creation and filtering

Using KEGG pathway information, pathways were grouped in hierarchical format (specific pathway (e.g., 00030 Pentose phosphate pathway) → Sub-category of Pathways (e.g., 1.1 Carbohydrate metabolism) → Category of Pathways (e.g., 1. Metabolism). Each category and sub-category of pathways contained information on name of the category, the number of pathways within the category, average direction of impact of pathways, average impact of pathways, the z-score of the flux and impacts, arithmetic means, standard deviation, sums of standard deviation for pathways, and an array of all pathway objects belonging to the group (e.g. ‘caffeine metabolism’ belongs in the group metabolism). The pathway class included the genes belonging to the pathway, the direction of impact of the gene on the pathway, fold-change of upregulated genes, fold change of downregulated genes, average *p*-values of upregulated genes, average *p*-values of downregulated genes, and proportion of DEG in the pathway corrected by FDR and minimum number of genes in background. The genes class contained the gene ID (Entrez gene ID), FDR, *p*-value for all comparisons, and fold change for all comparisons. Each grouping’s score (flux and impact) was derived from averages of the internal class (e.g. metabolism’s total flux is derived from the average of all metabolism pathways).

For each pathway, let N be the ratio of DEG that have an associated FDR of less than 0.05 (default value, can be changed) belonging to a specific pathway compared to the annotated genes in the background that belonging to that pathway and S be the minimum number of genes in the annotated background that are associated with a specific pathway. If N≤0.3 and S≤4 (can be changed depending on quality of data/reduction of outliers), the pathway was considered to have not enough information and the output was 0 for all comparisons.

### Step 3: Calculating flux and impact of a pathway

To separate the DEG, the FDR value was used (default = 0.05). The ratio of DEG in the pathway to annotated genes in the background associated with the same pathway was used in the calculation of impact. Let this ratio of DEG to background be R. Impact can then be expressed as

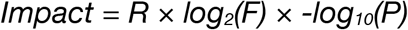

where F is the average absolute value of fold change for all the genes with an overall FDR≤0.05 in the pathway calculated, and P is the average *p*-value for the comparison of that time point with time point 0 for all the DEG (i.e., with an overall FDR≤0.05) in the pathway.

Flux (direction of impact) was then calculated as:

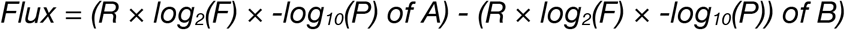

where A is up-regulated genes with a positive effect on the pathway and down-regulated genes with a negative effect on the pathway and B is down-regulated genes with a positive effect on the pathway and up-regulated genes with a negative effect on the pathway. A and B correspond to the assigned effect of each gene to each pathway following the pathway drawing provided by KEGG.

With sufficient annotation, the impact score as a signal for high expression activity and the flux score for the determination of directional magnitude (e.g. high expression activity but low-none flux may indicate balancing of a pathway) can be used to analyze time-course transcriptomic data.

### Step 4: Spline interpolation using R cubic spline function

Spline interpolation was calculated using R cubic spline function from the zoo.na library (Zeileis and Grothendieck, 2005). Because the cubic spline interpolation passes exactly through experimental points, key time points were preserved in the prediction and comparisons between those points still can be made. Each pathway for each dataset was splined using this function from time point 0 to the minimum time point among all datasets.

### Step 5: Curve comparison using z-normalization and auto/cross-correlation

Z-normalization was used to allow for comparisons between multiple experiments to establish trends. It can be performed because the values produced by pairwise comparisons are in relation to the control or time-point 0; thus each experimental time point is based on its statistical difference from the control. Normalization creates a parallel structure in which mean is approximately 0 and standard deviation is in a range close to 1. Z-normalization is also strong in structural pattern mining, as we were searching for structural similarities and dissimilarities, rather than amplitude variation.

Z-normalization was performed using an R programming language function as follows, using each pathway as an input vector and transformed into a resulting z-score output vector:

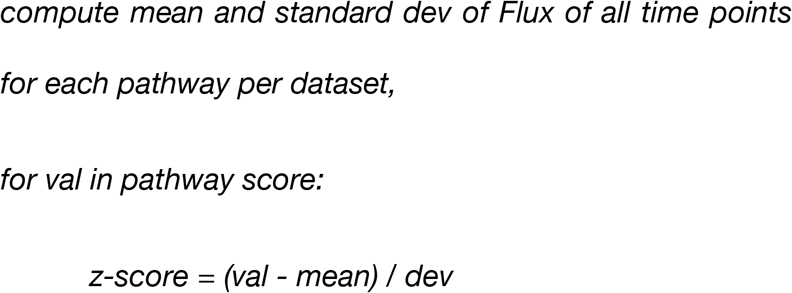

Normalization of the entire vector allowed for analysis between different experiments and identifying the peaks and valleys within the normalized distribution determines either outliers or pathway shape.

Cross correlation was performed after normalization of the matrices. For cross-species interpretation, each pathway was compared pairwise using R’s cross-correlation function ccf after normalization and a Pearson correlation coefficient was calculated. The threshold was calculated using standard error based on an uncorrelated series. A score above the threshold at 0 in the cross-correlation graph would indicate the best predictor is no time lag; a max score at any other x value would indicate that the x+k value is the best predictor. Given that r = 0.7 for the limit of experimental reproducibility, given a single species, and only 2-7% of genes overlap during stem cell differentiation (Evsikov and Solter, 2003), we used r≥0.6 as a benchmark for between time-lagged species correlation.

### DAVID Analysis

Using the large gene lists generated from the datasets, a sublist of genes with FDR≤0.05 was submitted to the Database for Annotation, Visualization and Integrated Discovery (DAVID) v6.8 (https://david.ncifcrf.gov/) for each dataset as the gene list. The total list of annotated genes was then submitted as the background [Huang et al 2008]. The resulting functional analysis chart and DAVID pathway viewer was then filtered by pathways with EASE score ≤0.05.

## Results and Discussion

### Implementation

The web tool DIA-norm was built using C++ and R. It is recommended that users use Chrome, Edge, or Firefox; however, it should work with any web browser. DIA-norm can currently be found at: http://bit.ly/osudiatool. The user first inputs a delimited file containing gene expression information. The file requires gene id (ENTREZ Gene ID), overall FDR of the effect of the condition studied, and, for each comparison, it requires fold change and *p*-value. The user then chooses the species of the dataset and can adjust any default parameters. These parameters include FDR cutoff, *p*-value cutoffs for individual comparison, the minimum number of genes measurable (i.e., background genes) in a pathway, and the minimum percentage of genes measurable in a pathway. Following the input phase, DIA-norm executes and displays a table where users can select pathways to display on the chart. DIA-norm offers 4 types of visualization: pathway impact, pathway direction, pathway group impact, and pathway group direction (Figure 1).

**Figure 1.**
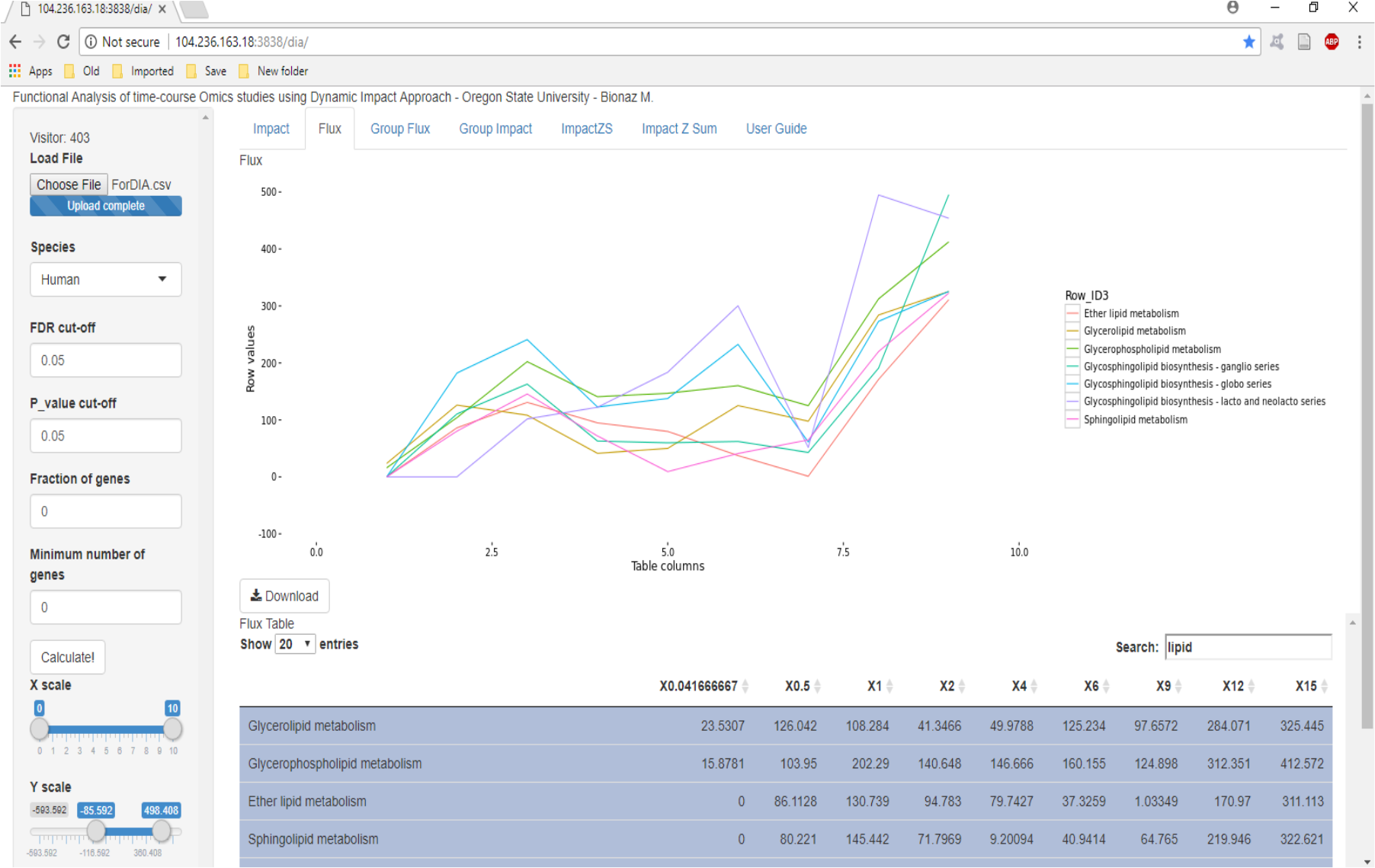
Example of DIA-norm flux calculation for selected pathways and time-points.

### Functional Similarities

DIA-norm has some functional similarities to other tools and methods. Steps 4-5 have similarities to the Bayesian network interpolation methods (e.g. dynamic time warping) (Bar-Joseph et al., 2002). Bar-Joseph et al. used a more refined method than the piecewise spline method used here, as their algorithm also took into account individual time alignment between different experiments. This approach is not applicable for DIA-norm, as DIA-norm’s functionality does not yet include simultaneous analysis of multiple datasets. Comparisons between species for DIA-norm is not yet automated, however the use of the individual time alignment would be the first step of extending the automation of the tool to evaluate the viability of a model organism.

### Analysis of Adipogenesis

The process of cell differentiation, specifically adipogenesis, was chosen for interspecies pathway analysis. Five experiments (microarray/next-gen seq) from NCBI GEO were chosen for their similarity in methods (3x human, mouse, pig). All experiments used mesenchymal stem cells that were differentiated *in vitro* toward adipogenesis and collected samples to run microarray analysis during several time points, including a baseline (or time 0).

DIA-norm has a key advantage compared to DIA is its ability to compare between species using normalization, interpolation, and cross-correlation analysis. This allows for pathways to be compared pairwise between species given similar KEGG pathway listings. Table 1 shows key pathways divided into four categories: early upregulation, middle upregulation, late upregulation, and middle downregulation. The full lists of these pathways can be found in supplementary files (DIAvsDavid.xlsx).

**Table 1.**
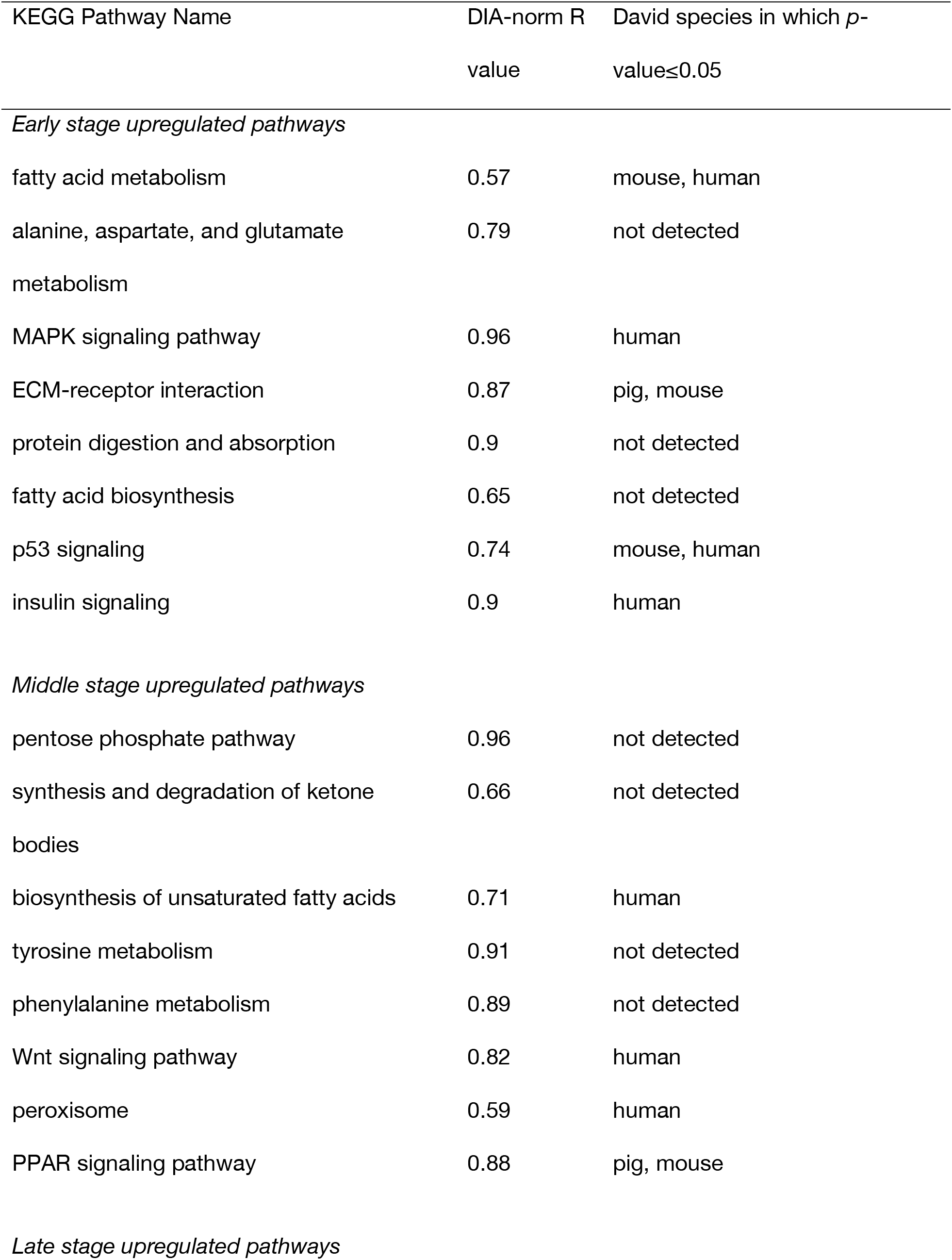
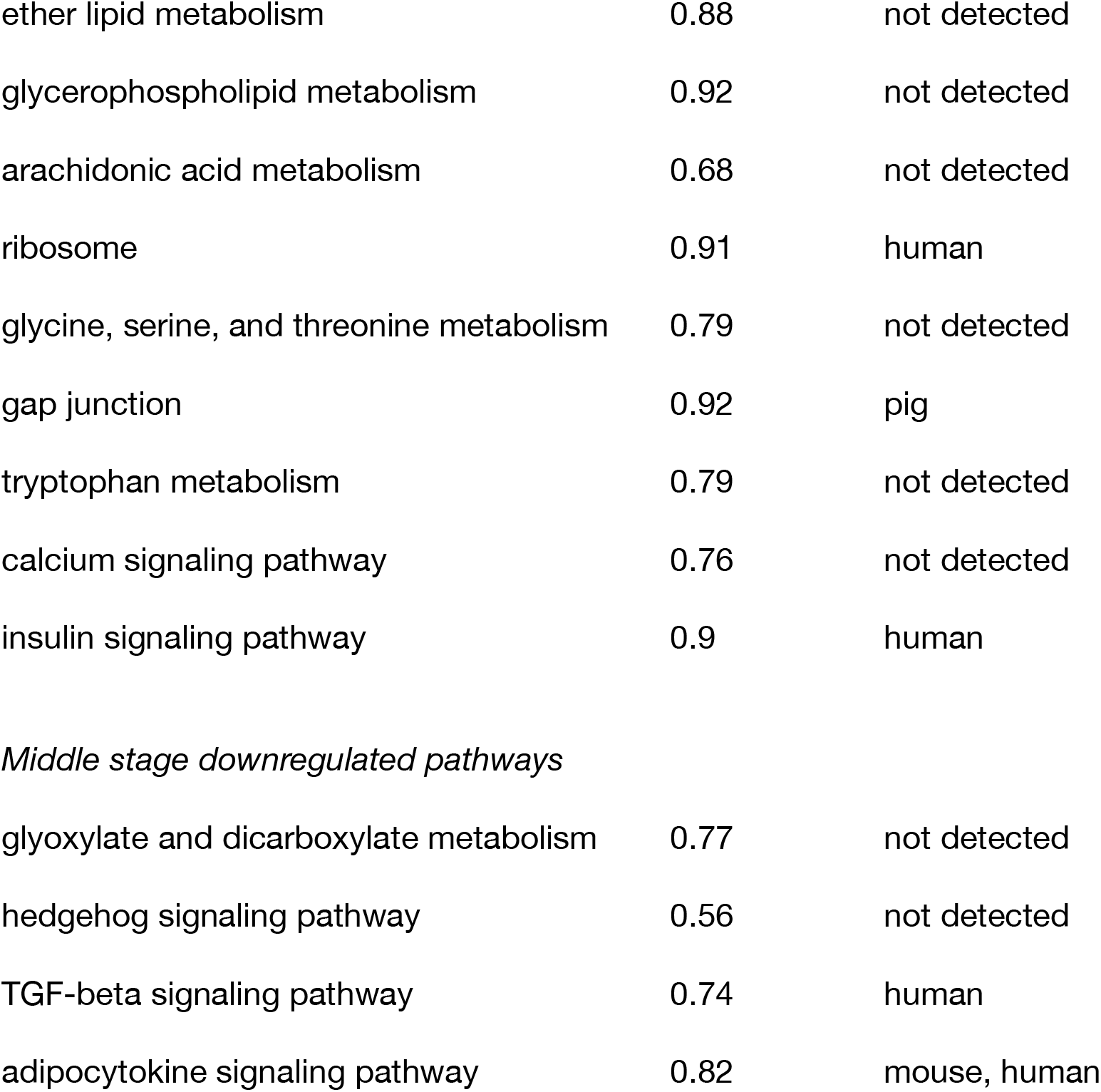
DIA-norm vs David for Adipogenesis-related Pathways

| KEGG Pathway Name | DIA-norm R value | David species in which $p$-value $\leq 0.05$ |
| --- | --- | --- |
| *Early stage upregulated pathways* |  |  |
| fatty acid metabolism | 0.57 | mouse, human |
| alanine, aspartate, and glutamate metabolism | 0.79 | not detected |
| MAPK signaling pathway | 0.96 | human |
| ECM-receptor interaction | 0.87 | pig, mouse |
| protein digestion and absorption | 0.9 | not detected |
| fatty acid biosynthesis | 0.65 | not detected |
| p53 signaling | 0.74 | mouse, human |
| insulin signaling | 0.9 | human |
| *Middle stage upregulated pathways* |  |  |
| pentose phosphate pathway | 0.96 | not detected |
| synthesis and degradation of ketone bodies | 0.66 | not detected |
| biosynthesis of unsaturated fatty acids | 0.71 | human |
| tyrosine metabolism | 0.91 | not detected |
| phenylalanine metabolism | 0.89 | not detected |
| Wnt signaling pathway | 0.82 | human |
| peroxisome | 0.59 | human |
| PPAR signaling pathway | 0.88 | pig, mouse |
| *Late stage upregulated pathways* |  |  |
| ether lipid metabolism | 0.88 | not detected |
| glycerophospholipid metabolism | 0.92 | not detected |
| arachidonic acid metabolism | 0.68 | not detected |
| ribosome | 0.91 | human |
| glycine, serine, and threonine metabolism | 0.79 | not detected |
| gap junction | 0.92 | pig |
| tryptophan metabolism | 0.79 | not detected |
| calcium signaling pathway | 0.76 | not detected |
| insulin signaling pathway | 0.9 | human |
| *Middle stage downregulated pathways* |  |  |
| glyoxylate and dicarboxylate metabolism | 0.77 | not detected |
| hedgehog signaling pathway | 0.56 | not detected |
| TGF-beta signaling pathway | 0.74 | human |
| adipocytokine signaling pathway | 0.82 | mouse, human |

The vast majority of adipogenesis-related pathways, in particular, the sugar and lipid metabolism pathways in which the five experiments showed correlation strongly supported existing literature by stage and direction. Not all correlated pathways from DIA-norm had information on stage and direction (e.g. leukocyte transendothelial migration) or had a consensus on the direction of impact (e.g. Wnt signaling pathway).

### Non-correlating Pathways

Glyoxylate and dicarboxylate have an interesting role in adipogenesis. In mice mitochondrial cells, oxodicarboxylate carrier is associated with adipogenesis (Das et all., 1999) (Niimi et al., 2009). However, DIA-norm results show that there is a negative valley in the middle stage (Figure 2a). There is a possibility that very early induction of this pathway drives this association (e.g. within the first 3 days). However, dicarboxylate carrier is expressed at an extremely high level throughout adipogenesis; thus it is unlikely to be the case (Das et al., 1999).

**Figure 2.**
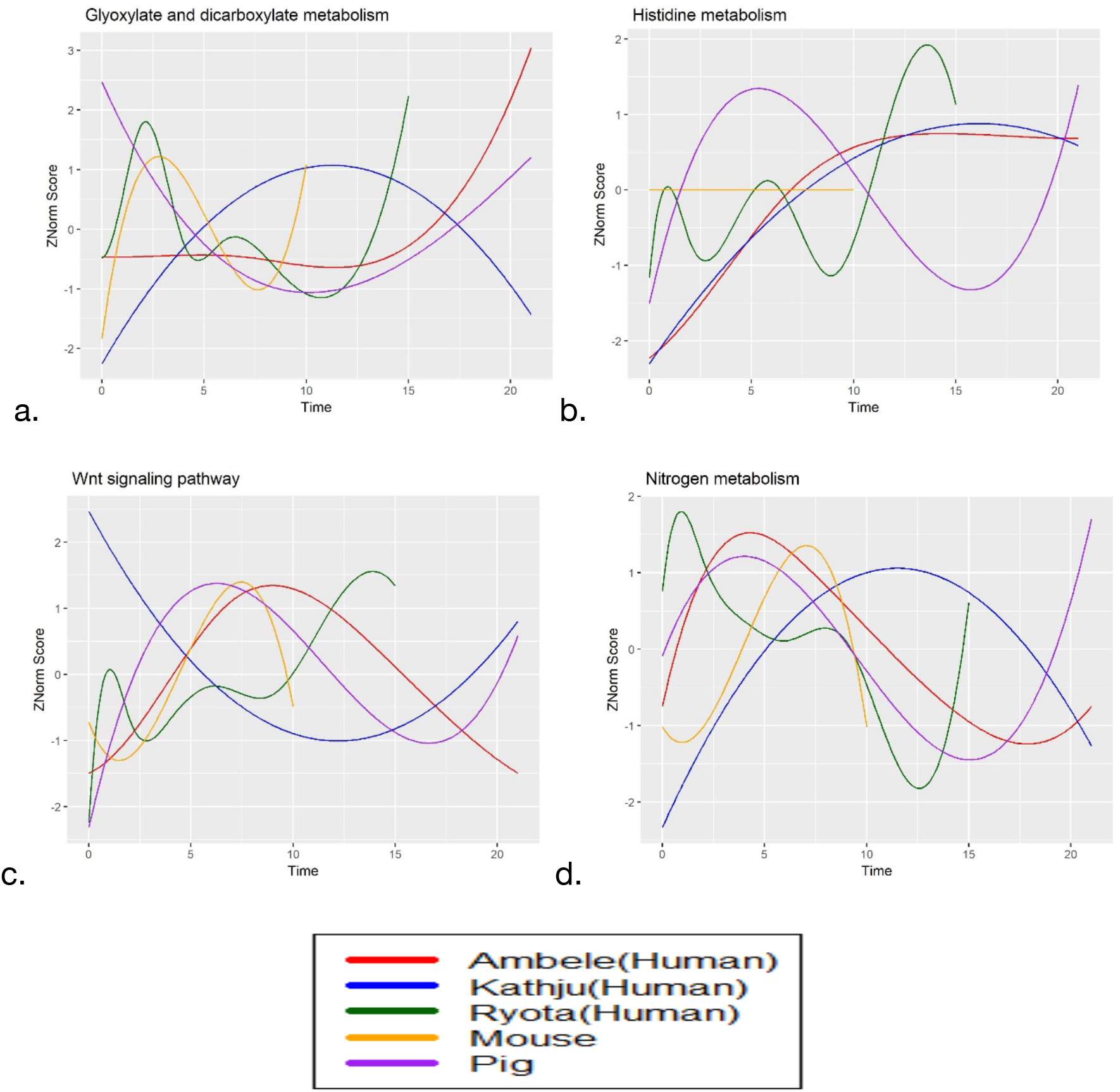
Expression of selected pathways using normalized scores versus time in days. The blue line (Satish et al., 2015) had the fewest data points (3), thus was noted as the least reliable for comparisons.

One of the more interesting pathways to examine is histidine metabolism. The effects of histidine in adipogenesis were contradictory to the literature (Ali et al., 2005, 2006a, 2006b, 2015; Aus Tarig, 2004). Histidine acts as an ALP (alkaline phosphatase) inhibitor and as an adipogenesis downregulator according to the studies. The KEGG pathway diagram shows a positive relationship between the histidine metabolism genes and L-histidine. The results from DIA-norm showed an upward expression of the histidine metabolism pathway in 4 of the 5 experiments. If histidine is a downregulator of adipogenesis, the histidine metabolism pathway should decrease during the early to middle stages of adipogenesis. In all the studies conducted at the University of Witwatersrand (Ali et al., 2005, 2006a, 2006b, 2015; Aus Tarig, 2004), human preadipocytes and 3T3-L1 cells were used, which are the same as the cells used in the DIA-norm analysis. All of the information on histidine’s role in adipogenesis comes from this single research group.

Interestingly, the signaling pathways exhibit more variation via higher z-scores than metabolic pathways. One example of a signaling pathway showing contradictory information is Wnt signaling (Figure 2c). Wnt signaling suppression is part of adipogenesis, but the same research offers some caveats (Christodoulides et al., 2009; Wolfson et al., 2015). Wnt5b, for example, is transiently induced during middle stage adipogenesis and acts to destabilize beta-catenin and promotes further differentiation, whereas other Wnt molecules such as Wnt10b stabilizes beta-catenin and represses differentiation (Moldes et al., 2003). It is theorized that PPARγ plays a significant role in deactivating Wnt10b and activating Wnt5b, thus suppressing the mechanism that maintains the undifferentiated states of preadipocytes (Prestwich and Macdougald, 2007).

Wnt signaling pathway’s inhibiting effect on adipogenesis would suggest that it should be downregulated during adipogenesis in order to break down beta-catenin and allow for differentiation to occur. Four of the five experiments tested show induction of Wnt signaling with at least one peak in early-middle stage. The other experiment that did not show the same pattern (human experiment 3) presented the least number of time points, and thus, interpolation is less robust. This difference in regulation may suggest that Wnt10b related genes were not included in the expression profile or were not significant enough to make the 0.05 adjusted cut-off. Looking at the genes in the Wnt pathway in the KEGG database, cell adhesion, MAPK, and p53 signaling pathway genes are incorporated within the Wnt pathway, all of which are upregulated during adipogenesis. Thus, there is a possibility that Wnt10b related genes may have been marginalized by the large number and high fold changes of other signaling genes such as MAP3K7 or NLK.

### Correlating Pathways

Among all pathways, non-cancer disease pathways (immune, neurodegenerative, cardiovascular, metabolic, infectious) had the highest correlation. For this analysis, we used r≥0.6 (*p*-value*≤0.05)*. Specifically, when the disease type was heart and epithelial tissue related (e.g. viral myocarditis), correlation coefficients reached above 0.8. We found that the overall order of correlation for all pathways of species from largest to smallest was: mouse-pig (0.57), human-pig (0.51), human-mouse (0.46). Surprisingly, metabolism had fairly low correlation percentage (Table 2). However, in specifically adipogenesis and fat related pathways, the correlation (Table 1, Supplementary file DetailedPathwayCategories.xlsx) was much higher.

**Table 2.** Breakdown of Pathways via KEGG Pathway Hierarchy

| Pathway Type (KEGG A/B Level) | Number of Pathways | Number of correlating pathways, $r \geq 0.6$ | Correlation Percentage, correlating/total pathways in category |
| --- | --- | --- | --- |
| Metabolism | 86 | 36 | 41.9 |
| Environmental Information Processing | 50 | 31 | 62.0 |
| Cancer | 15 | 6 | 40.0 |
| Other Disease Pathways (Immune, Neurodegenerative, Cardiovascular, Metabolic, Infectious) | 30 | 25 | 83.3 |

Nitrogen metabolism (Figure 2d) resulted in unusually high correlation (r = 0.86). In fact, nitrogen metabolism had the highest correlation values for any metabolic pathway. There are currently very few studies linking the role of nitrogen metabolism to adipogenesis. However, its link to glutamate may play an important role in regulating adipogenesis (Green et al., 2016). According to the KEGG pathway diagrams, nitrogen metabolism can create the metabolite L-glutamate, which has extremely high uptake and synthesis in the initial stages of adipogenesis.

### Model Determination

The pig dataset had a higher number of correlating pathways and higher average correlation score (64.5 average pathways, r = 0.51) with human than the mouse (30.5 average pathways, r = 0.46) and had data points up to 21 days, whereas the mouse dataset had points up to 10 days (mouse matures relatively quickly compared to human/pig). There is a possibility that the mouse data would have been more highly correlated had there been data points past 10 days. Thus, we cannot definitively conclude that a pig would act as a better model than the mouse; however, the DIA-norm findings are in accordance with prior knowledge that the pigs are a good model for studying humans, specifically regarding obesity (Koopmans and Schuurman, 2015). The average R value of the two human experiments at the n>3 time points, the average R value was fairly low (0.49), compared to the pig vs mouse (0.57) and human vs pig (0.52). For metabolic and disease pathways specifically, however, the human values (0.52 metabolic, 0.56 non-cancer disease) averaged to be greater than human vs combined mouse/pig model (0.48 metabolic, 0.54 non-cancer disease).

### Comparisons

We compared the results of DIA-norm to a functional annotation tool, DAVID, and found that DAVID missed a large number of significant pathways (Tables 1,3) (Supplementary file DIAvsDavid.xlsx). For example, DAVID misses glycerophospholipid metabolism and hedgehog signaling pathways entirely, where they should show significance due to their roles in regulating adipogenesis (Fleury et al., 2016; Fontaine et al., 2008; Hishikawa et al., 2014).

**Table 3.** DIA-norm vs DAVID Total Pathways Detected Comparison (includes non-significant pathways)

| Experiment | Number of<br>DAVID<br>pathways<br>detected | Number of<br>DAVID<br>pathways $p$-<br>value≤0.05 | Number of<br>DIA-norm<br>pathways<br>detected | Difference in total pathways<br>captured, DAVID vs DIA-norm |
| --- | --- | --- | --- | --- |
| Human1 (Kathju) | 138 | 120 | 221 | -83 |
| Human2 (Ryota) | 114 | 97 | 211 | -67 |
| Mouse | 34 | 24 | 167 | -133 |
| Pig | 6 | 4 | 34 | -28 |

The mouse dataset only had 24 pathways that had a large enough number of genes to result in an overall *p*-value≤0.05, missing biologically important pathways such the nucleotide metabolism pathways, cAMP and Wnt signaling, and many of the lipid metabolism pathways. Even smaller is the number of significant pathways discovered in the pig dataset. Only 4 of the 6 identified pathways resulted in a *p*-value of ≤0.05 and only found 3 biological relevant pathways: gap junction, ECM receptor interaction, and ppar signaling pathway. While these pathways, specifically ppar signaling, are important for adipogenesis, many other important pathways (Table 1) were not detected as significant.

## Limitations

Functionally, DIA-norm does not use an unsupervised reduction of gene lists that many other tools implement, and thus allows for a holistic view of pathways through time; however, it has its own series of limitations. One of the main limitations is the weights of the genes. Without a clustering approach, highly connected genes are weighted the same as any other gene; thus a pathway with genes that have large expression values but are biologically less significant can produce higher scores than a pathway with lower expression values but biologically more significant affected genes. Currently, there are efforts to provide weights to genes, although these works are incomplete and do not include biological validation studies (Du et al., 2016).

For the analysis of adipogenesis, drawing conclusions and evaluating validity of a model organism from the data are limited because not all the experiments used the same adipogenic cocktail and not all the experiments used the same type of cells. The mouse dataset was collected from a pre-adipocytes cell line (3T3-L1) whereas the human and pig were mesenchymal stem cells collected from different individuals. This can partly explain the lower correlation observed with human and human and the higher correlation detected between pig and human. The best model would have been the use of primary mesenchymal stem cells for mouse or also a comparison between mouse cell line and human cell line, such as SGBS. One counter to this limitation is that the pig vs mouse correlation was very strong regardless of type of cells; however, cells from pigs and mice underwent almost identical similar adipogenic induction. This suggests that other phylogenic factors led to the pig being the stronger animal model for adipogenesis. This is validated by general studies on the validity of the pig model in human disease treatment and further supported by the phylogenic research (Wernersson et al., 2005).

## Conclusion

We have introduced DIA-norm, a standalone software tool for the functional analysis of time-series datasets for human, mouse, pig, and cow. The software found biologically relevant pathways, identified direction of biological impact and its magnitude, and compared patterns of pathways between species. It presents its information in an interactive visualization of line graphs designed for time-series data. DIA-norm’s integration with KEGG and gene direction databases allowed for easy biological interpretation, and as those databases are curated and become more populated, the capabilities of the tool will grow in sync. DIA-norm is designed for time-series data and analyzes each comparison independently. When the z-score is calculated for comparison between species, it makes the assumption that the data are sequentially ordered. Thus, it is also capable of analysis of sequential experiments such as treatment dosage response.

DIA-norm identified the vast majority of biological important pathways. For the process of adipogenesis, it achieved 80+% pathway correlation of well-studied pathways relating to adipogenesis at a high correlation cut-off. DIA-norm can be used for both time-series pathway analysis and the determination of a model organism. Despite the limitations as discussed above, DIA-norm predicted that the pig is a better model to humans than a mouse for the study of adipogenesis. Our findings indicate that DIA-norm can be used to study the effect of any treatment, including drugs, on specific pathways between multiple species to determine the best animal model for that treatment for future studies. The reliability of DIA-norm to provide biological insights compared to enrichment approach tools has been demonstrated in the selected transcriptomic studies by identifying a higher number of total and biologically relevant pathways. DIA-norm’s final advantage is its highly graphical outputs that are easy to interpret for both biologists and the public audience.

